# Limited genomic signatures of population collapse in the critically endangered black abalone (*Haliotis cracherodii*)

**DOI:** 10.1101/2024.01.26.577275

**Authors:** Brock Wooldridge, Chloé Orland, Erik Enbody, Merly Escalona, Cade Mirchandani, Russell Corbett-Detig, Joshua D. Kapp, Nathaniel Fletcher, Karah Ammann, Peter Raimondi, Beth Shapiro

## Abstract

The black abalone, *Haliotis cracherodii*, is a large, long-lived marine mollusc that inhabits rocky intertidal habitats along the coast of California and Mexico. In 1985, populations were impacted by a bacterial disease known as withering syndrome (WS) that wiped out >90% of individuals, leading to the species’ designation as critically endangered. Current conservation strategies include restoring diminished populations by translocating healthy individuals. However, population collapse on this scale may have dramatically lowered genetic diversity and strengthened geographic differentiation, making translocation-based recovery contentious. Additionally, the current prevalence of WS is unknown. To address these uncertainties, we sequenced and analyzed the genomes of 133 black abalone individuals from across their present range. We observed no spatial genetic structure among black abalone, with the exception of a single chromosomal inversion that increases in frequency with latitude. Genetic divergence between sites is minimal, and does not scale with either geographic distance or environmental dissimilarity. Genetic diversity appears uniformly high across the range. Despite this, however, demographic inference confirms a severe population bottleneck beginning around the time of WS onset, highlighting the temporal offset that may occur between a population collapse and its potential impact on genetic diversity. Finally, we find the bacterial agent of WS is equally present across the sampled range, but only in 10% of individuals. The lack of genetic structure, uniform diversity, and prevalence of WS bacteria indicates that translocation could be a valid and low-risk means of population restoration for black abalone species’ recovery.

## 1 Introduction

Severe population declines threaten a species’ long-term viability and can even result in extinction. Although conservation of remnant populations is essential to maintain any hope of recovery, a key question is whether the genetic effects of such a decline (i.e. bottleneck) are too deleterious to overcome (Robinson et al., 2022). Smaller populations are expected to experience inbreeding depression, or reduced fitness as a result of increased mating between related individuals (D. Charlesworth & Willis, 2009; Keller & Waller, 2002). Natural selection is sometimes thought to be less effective at removing mildly deleterious alleles in small populations, permitting the accumulation of deleterious variation (i.e. mutation load) (Agrawal & Whitlock, 2012; Lynch et al., 1995). Finally, reduced overall variation in smaller populations is also thought to limit the potential to adapt to new environments (B. Charlesworth, 2009; Frankham et al., 1999; Hoffmann et al., 2017). While the relationship between these phenomena, genetic diversity, and extinction risk is complex (Kardos et al., 2021; Teixeira & Huber, 2021), the field of conservation genomics continues to play an essential role in guiding the recovery of small populations (Shaffer et al., 2022).

Genomic data can provide surprising insights into the status of small populations. Recent work has revealed unexpected viability of critically endangered species, providing evidence that just 10 individuals of the vaquita porpoise may be sufficient to avoid inbreeding depression and seed recovery (Robinson et al., 2022). Similarly, genomic data have shown that long-term small population size has actually enabled Channel Islands foxes to effectively purge highly deleterious variants, suggesting that genetic rescue through introduction of new individuals may not be an appropriate strategy (Robinson et al., 2018). Even when expected, the outcomes of conservation genomics studies can still provide valuable guidance for species management. Analyses of the Alpine ibex confirmed the persistent genomic signature of near-extinction despite a healthy current population size, providing thresholds for minimum population size to avoid inbreeding depression (Grossen et al., 2020). As evidenced by these cases, genomic data can provide a helpful lens to understanding the trajectory of a threatened population (Willi et al., 2022). With genomic resources becoming increasingly available for non-model organisms, including those that were once prohibitively difficult to access and sequence, it becomes imperative to integrate this information into management and recovery strategies.

Black abalone (*Haliotis cracherodii,* Leach 1814) are large, long-lived marine gastropods found along roughly 1,500 km of the coastline from Point Arena, in California, USA, to Bahia Tortugas and Isla Guadalupe, in Baja California, Mexico (Neuman et al., 2010). They typically live in rocky intertidal habitats and less often subtidally to a depth of 6 meters, and occupy a key niche in intertidal ecosystems as primary consumers of macroalgae (Leighton & Boolootian, 1963) and common prey items for sea otters (Raimondi et al., 2015). Black abalone facilitate encrusting coralline algae, thereby maintaining favorable habitat for conspecific recruitment on rocky intertidal reefs (Cox, 1962; Miner et al., 2006; Richards & Davis, 1993). Black abalone are dioecious and reproduce by broadcast spawning. While this reproductive strategy may facilitate gene flow between distant populations, the negative buoyancy of embryos and the 5-15 day larval swimming phase that follows is thought to limit dispersal in comparison to other broadcast spawners (Chambers et al., 2006).

In addition to playing a key ecosystem role, the abundance, accessibility, and size of black abalone have made them a frequent target of human populations for at least 13,000 years (Haas et al., 2019). The meaty foot has served as a food staple for indigenous Californians, and their iridescent shells have been used as adornments, tools, cultural currency, and religious symbols (Erlandson et al., 2008; Kelley & Francis, 2003; Sloan, 2003; Vileisis, 2020). After colonists from Spain and the United States replaced indigenous peoples through displacement, disease, and violence from the 17th-19th centuries, abalone were thereafter heavily impacted by commercial fishing (Bentz & Braje, 2017; Braje et al., 2014). These fisheries produced catches that are inconceivable today; in 1973 alone around 800 tons of black abalone were harvested from the California Channel Islands (Karpov et al., 2000).

Despite this species being arguably the hardiest of the seven abalone species found along this coastline (Tissot, 1988; Vileisis, 2020), black abalone were nearly erased during the late 20^th^ century (Rogers-Bennett, 2002). While intensified fishing and environmental pressures (e.g. oil spills, sea temperature rise, sediment burials) contributed to this decline, the primary culprit was the emergence of a devastating disease known as withering syndrome (WS). WS is caused by the Rickettsia-like bacteria *Candidatus Xenohaliotis californiensis*, also known as WS-RLO, which attacks the lining of the digestive tract, resulting in reduced body mass and eventual withering of the abalone’s foot until it can no longer cling to the substrate (Friedman et al., 2000; Lafferty & Kuris, 1993). Following the onset of WS around 1985, black abalone underwent widespread mass mortality events. In areas most affected by the disease, populations declined by up to 99% (Crosson et al., 2014; Neuman et al., 2010; VanBlaricom et al., 2009). These dramatic declines led to the closure of the 150-year old fishery in 1993, and in 2009 the species was listed as endangered under the U.S. Endangered Species Act. Changes to intertidal ecosystems across the range followed this collapse, with habitats previously dominated by crustose coralline algae and bare rock becoming overgrown with fleshy algae and sessile invertebrates (Miner et al., 2006).

WS has been most severe in populations south of California’s Point Conception, including the California Channel Islands, and appears to be exacerbated in populations experiencing anomalously warm water (Ben-Horin et al., 2013; Crosson & Friedman, 2018), following warm El Niño cycles (Raimondi et al., 2002) or in the vicinity of power plant outflows (Altstatt et al., 1996). However, the mechanisms governing susceptibility are largely unresolved, in large part due to the overwhelming severity of the disease and the lack of suitable populations for study. The recent limited recovery of some populations indicates that WS immunity may exist, and this immunity has been tenuously linked to both a) a phage that infects the WS-causing bacteria and b) heritable genetic variation. Observations of a phage hyperparasite, known as *Xenohaliotis* or RLOv, infecting the bacteria led to the discovery that phage presence appears to partially attenuate WS development, but like the primary abalone-Rickettsia relationship this effect is also dependent on temperature (Friedman et al., 2014). At the same time, some populations - and other species - appear to have evolved limited resistance to WS even in the absence of phage (Brokordt et al., 2017; Crosson et al., 2014). However, evidence for both phenomena is preliminary, and it is unclear to what extent black abalone across the range have adapted to the spread of WS.

It is against this backdrop of overfishing, disease spread, and rapid decline that a genetic approach to species conservation has become particularly needed. Although substantial effort has gone towards understanding the status of remaining populations, little has been done through the lens of conservation genomics (Gruenthal & Burton, 2008; Hamm & Burton, 2000). Genomic data from wild black abalone can answer essential questions about remaining genetic diversity, connectivity between sites, and genetic associations with population success, that together with decades of ecological research will help guide critical management decisions. To this end, we developed a low impact external swabbing method for obtaining whole-genome data from wild black abalone, and sequenced 133 individuals from across ∼800 km of the former range. With these data we present the most comprehensive picture yet of population structure in this critically endangered species. Additionally, we explore how patterns of gene flow and genetic diversity associate with geographic and environmental variation to assess the extent to which the few remaining populations are genetically isolated. Finally, we determine the distribution of the bacterial agent of withering syndrome and explore associations with its phage hyperparasite.

## 2 Results

### 2.1 Whole genome sequencing of black abalone via DNA swabs

Because black abalone adhere strongly to their substrate, injuries during removal are often fatal, and obtaining substantial tissue clips risks serious harm to the animal. To avoid this risk, we tested whether high quality whole genomes could be obtained from swabbing the exposed edge of an individual’s foot with a sterile nylon-tipped swab. Using this approach, we swabbed 150 healthy abalone across 35 sites spanning the black abalone range (Fig. 1) and obtained high quality DNA extracts from 133. Average endogenous content was 63% abalone DNA, with significant proportions of reads mapping to each individual’s internal and external microbiome, including known parasites and symbionts (see section 3.5). Libraries were sufficiently complex to generate 5-20-fold genomic coverage per individual.

**Figure 1.**
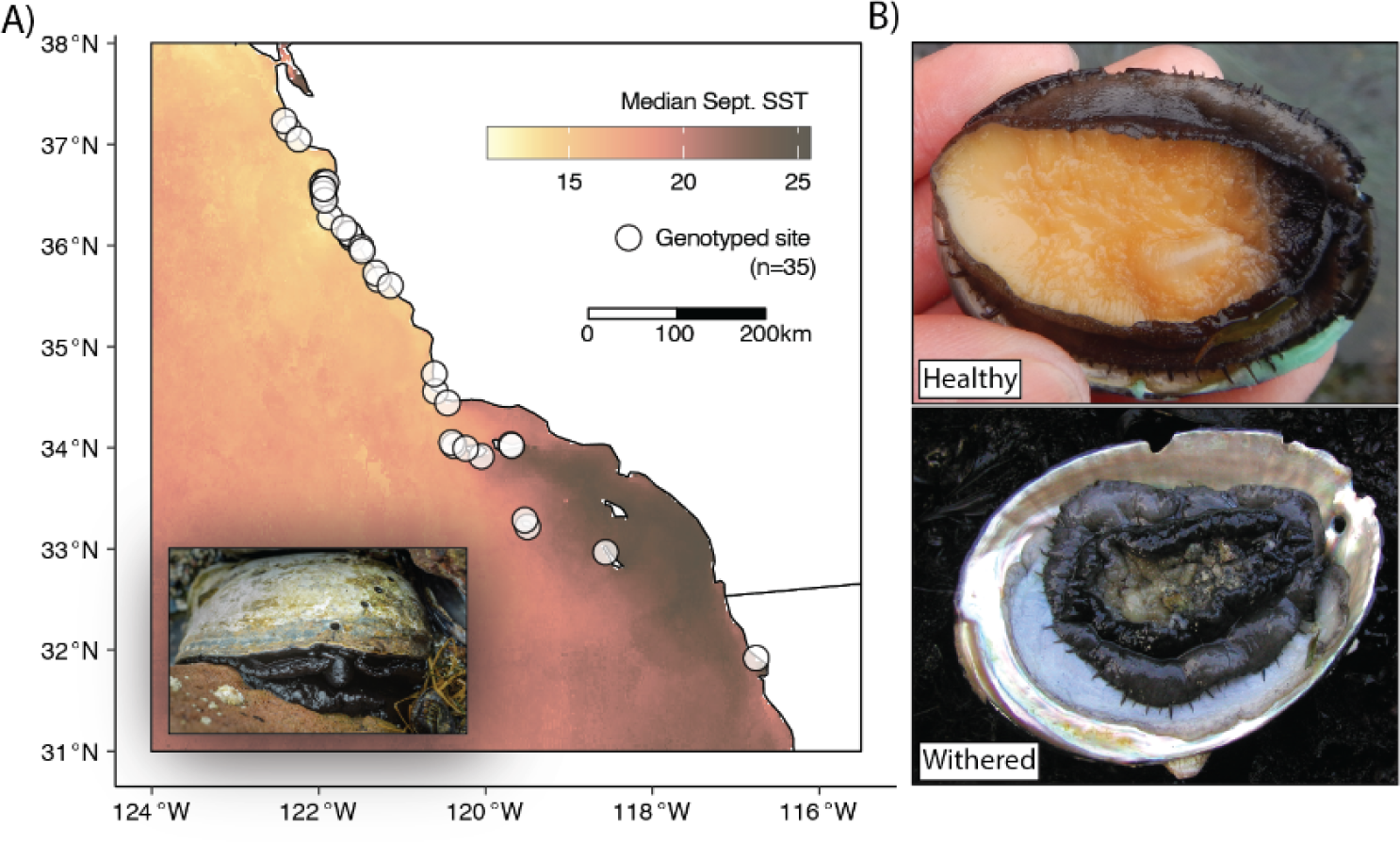
**A)** Distribution of collection sites along the California and Baja California coast. Inset displays adult black abalone, image by © Michael Ready. **B)** Representative images of healthy and withered black abalone. Photos by Nathaniel Fletcher.

### 2.2 Population structure along the California coast

A principal components analysis of unlinked genome-wide SNPs showed all samples clustering into one of three discrete groups (Fig. 2A). However, the principal component associated with this clustering (PC1) explained only 2.0% of the variation in this data. To understand what may be driving this pattern, we performed local sliding-window PCA with *lostruct* (Li & Ralph, 2019) and identified a large 31MB region on chromosome 4 significantly impacting genome-wide population structure (Fig. 2D). Interestingly, a PCA of SNPs from this region (Fig. 2B) showed identical group membership to the whole-genome PCA, suggesting that population structure at this locus alone is driving the whole-genome picture. When we excluded this region, which comprises 2.6% of the genome, from our analysis, this three-group structure disappeared from the PCA, resulting in a cloud of samples with little discernible structure (Fig. 2C).

**Figure 2.**
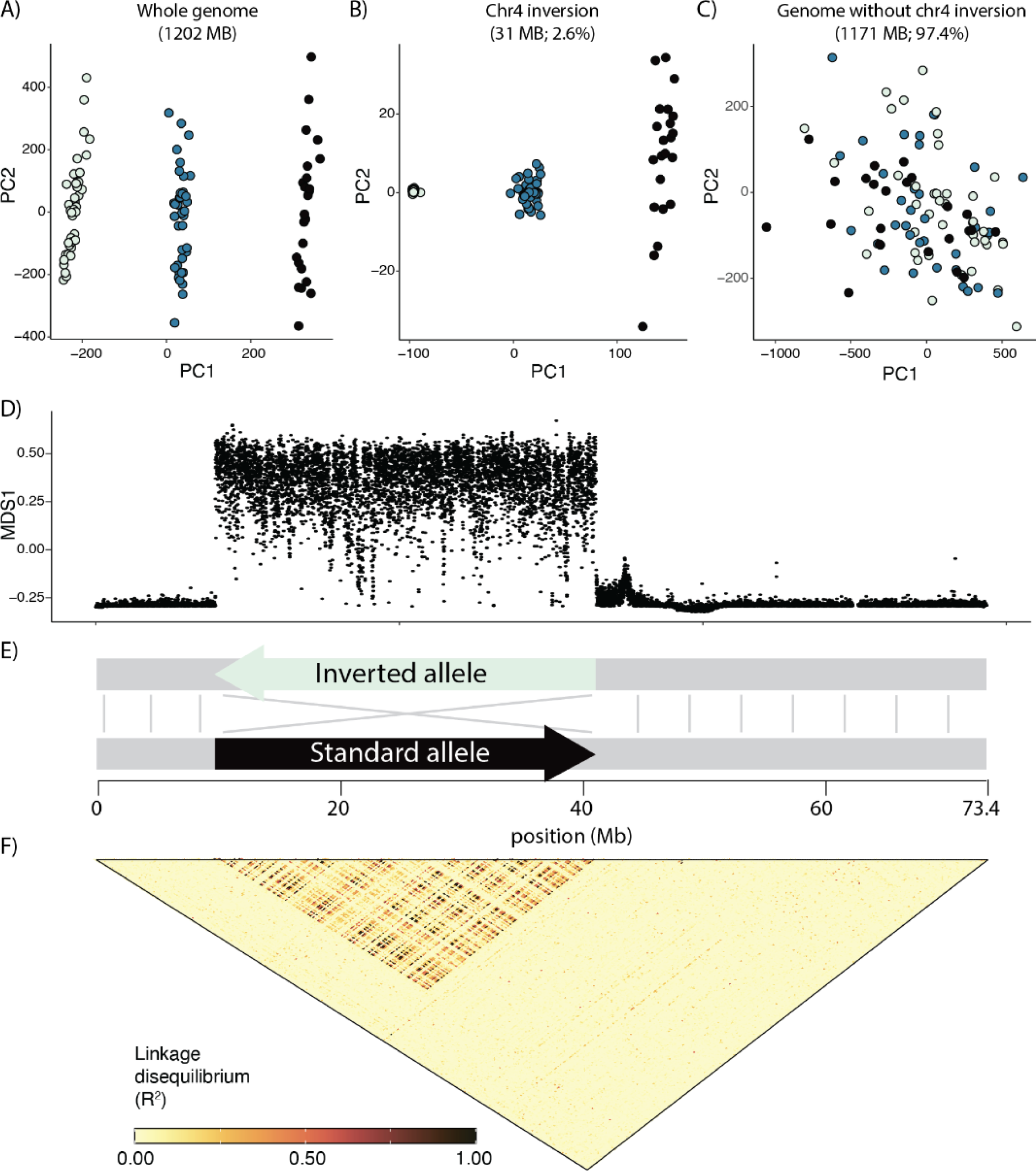
**A)** Genetic PCA based on unlinked genome-wide SNPs. In this panel colors are arbitrarily assigned to the three clusters. **B)** Genetic PCA based on 1kb-separated SNPs from the chromosomal inversion. Colors here correspond to those initially assigned in the genome-wide PCA (left). **C)** Genetic PCA based on unlinked genome-wide SNPs, excluding the chr4 inversion. Colors here correspond to those initially assigned in the genome-wide PCA (far left). **D)** Sliding-window PCA (*lostruct*), each point corresponds to the local genomic structure in 5000 bp genomic windows, with the “MDS1” position roughly representing deviation in the window from genome-wide structure. **E)** Simplified schematic showing size and structure of chr4 inversion. **F)** Linkage disequilibrium across chr4.

Further analysis of this anomalous region revealed a large chromosomal inversion (Fig. 2E). A Hi-C contact map based on the diploid reference genome (Orland et al., 2022) showed two potential scaffold arrangements directly corresponding to the distinct *lostruct* peak at chr4: 9.8 - 41.2 Mb (Fig. S1A). These two alternative arrangements map in opposite directions to each other, suggesting that this reference genome is heterozygous for a chromosomal inversion. Long-read variants also point towards an inversion in this region, although the suggested breakpoints only roughly correspond to those hinted at by the prior analyses (Fig. S1B). Finally, linkage-disequilibrium analysis of all 133 sequenced individuals showed elevated LD as might be expected if recombination-suppressing inversions were present (Fig. 2F; Hager et al. 2022). Together, these lines of evidence suggest that a 31MB non-recombining inversion on chr4 is underlying the three distinct clusters in the whole-genome PCA. This inversion is polymorphic in our samples (allele frequency = 58%); while the ancestral state is not known, the severely reduced polymorphism in the left homozygous group (Fig. 2B; Fig. S3) is suggestive of a recent origin and possibly selection, leading us to designate that allele as “inverted” and the alternate allele as “standard” (Fig. 2E).

### 2.3 Isolation-by-distance (IBD) and isolation-by-environment (IBE)

Genetic divergence between collection sites was low, with Hudson’s F_ST_ ranging from 0.02 to 0.07 (excluding chr4 inversion). This limited differentiation across the range agrees with the pattern in the “no-inversion” PCA (Fig. 2C) suggesting little overall genetic structure. However, even limited genetic divergence can still encompass important locally adaptive variation. Therefore, we aimed to assess what physical or environmental factors, if any, may be associated with genetic structure along this range.

We observed little relationship between the extent of genetic divergence and physical or oceanographic distance between sites, known as isolation-by-distance (IBD; Wright 1943). The relationship between F_ST_ and physical distance was weak but significant, with the linear model showing F_ST_ increasing by only 0.00035 for every 500 km of distance between sites (R^2^ = 0.01, p < 0.001; Fig 3A). Because black abalone are a marine species and ocean currents on the west coast of North America can be asymmetric, we then modeled the relationship between genetic distance and ‘connectedness’ between sites; that is, the probability of larvae dispersing (PLD) from one site to the next at 5, 10, and 15 days. Even for the 15 day model, which is roughly half the absolute maximum larval duration of black abalone (Morse et al., 1979), we observed almost no relationship between genetic distance and the probability of larval dispersal between sites (north to south: R^2^ = 0.02, p < 0.001; south to north: R^2^ = 0.01, p = 0.279; Fig. 3B).

**Figure 3.**
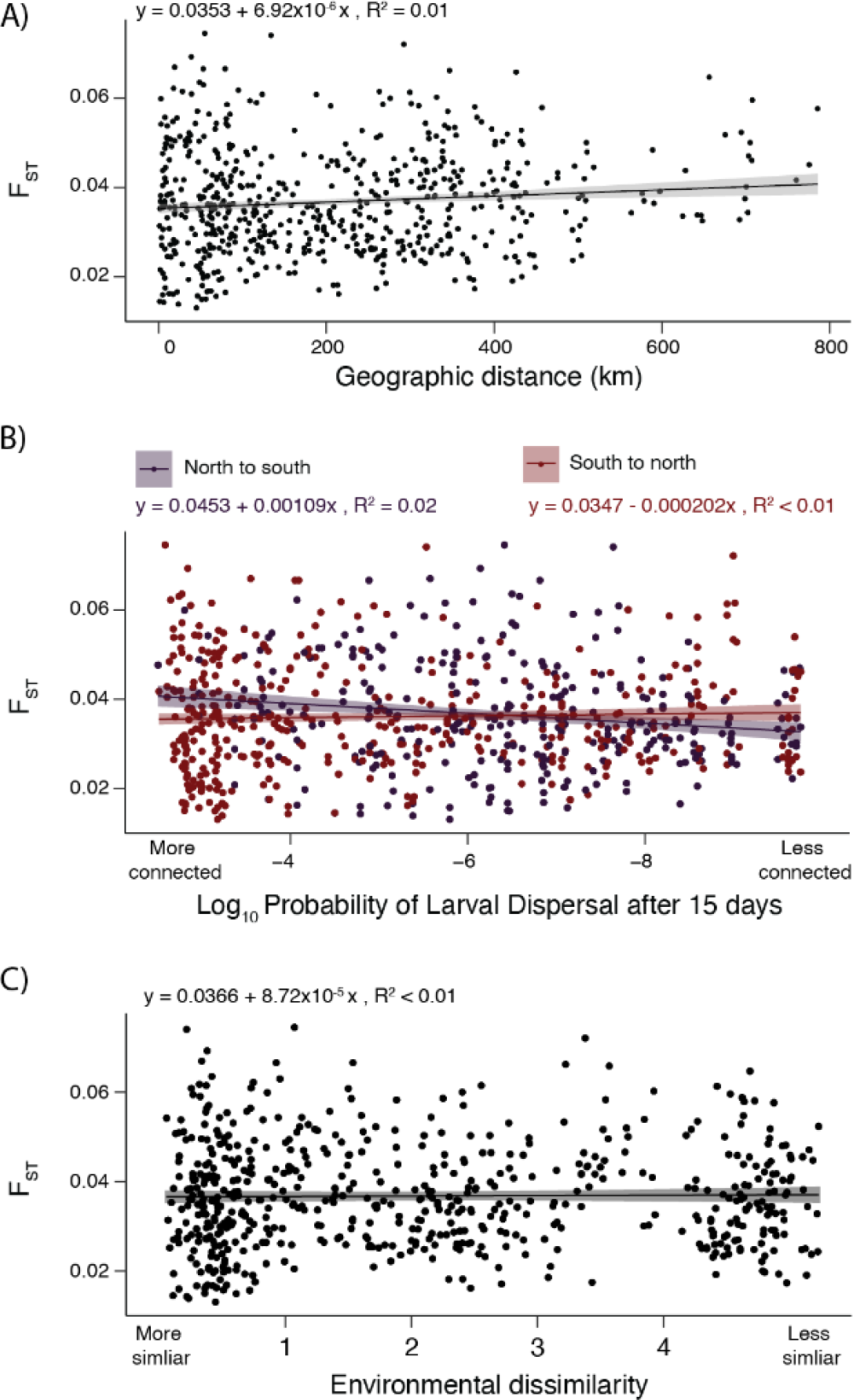
**A)** Isolation-by-Distance: genetic divergence between sites (F_ST_) as a function of straight-line distance between sites. **B)** Isolation-by-Distance: F_ST_ as a function of the probability of pelagic larvae dispersing (PLD) between sites within 15 days. Results for north-to-south and south-to-north are plotted. **C)** Isolation-by-Environment: F_ST_ as a function of environmental dissimilarity between sites. Environmental dissimilarity encompasses air temperature, sea temperature, and pH.

We also found no relationship between the extent of genetic divergence between abalone from any two collection sites and the similarity of their environments, also known as isolation-by-environment (IBE; Wang & Bradburd, 2014). To quantify environmental distance between collection sites, we measured differences in key environmental variables known to affect black abalone fitness. We summarized differences in air temperature, water temperature, and pH between collection sites via principal components analysis (Fig. S2). We found no association between the extent of environmental mismatch between sites - as quantified by PC1 - and genetic distance (R^2^ = 0.01, p = 0.671; Fig. 3C).Therefore, sites with more similar conditions, for example sea surface temperature, are not necessarily more closely related than those at locations with dissimilar conditions.

In contrast to the genome-wide patterns described above, we observed a striking north-south cline in the distribution of the chr4 inversion. The linear cline fit to the frequency of the inversion allele is steeper than 100% of linear clines fit to 14.9 million randomly selected SNPs (Fig. 4A). Indeed, the frequency of the inversion is significantly associated with latitude, with the inverted allele being more common in the north (Figs. 4 B,C; mean GWAS p-val = 1.59e-08). Therefore, while variation in the majority of the black abalone genome shows no segregation according to geography or environment (Fig. 3), the distribution of the chr4 inversion, encompassing 2.6% of the genome, is one clear exception to this general pattern.

**Figure 4.**
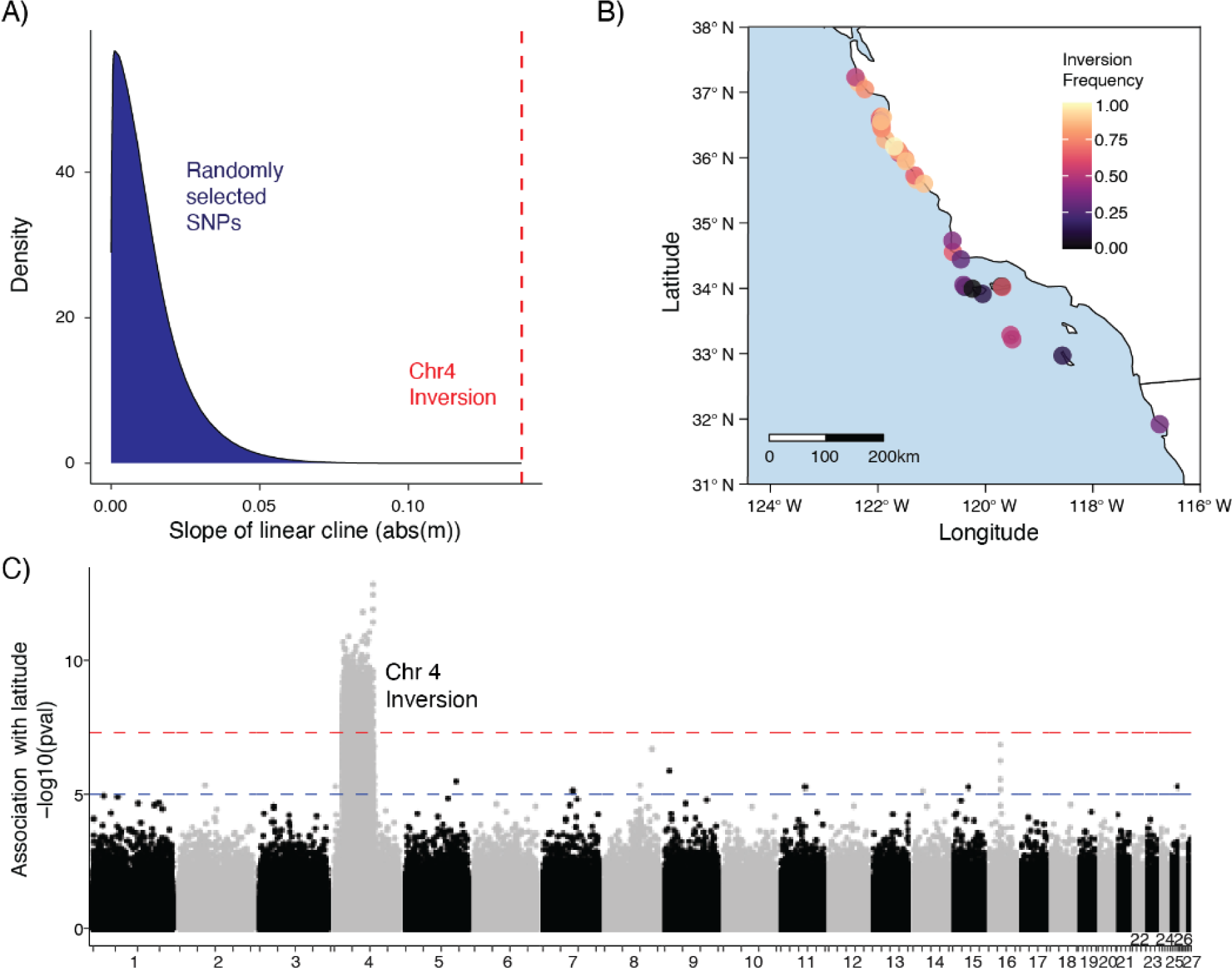
**A)** Distribution of coefficients for linear clines fit to site frequencies for ∼14e6 SNPs, as compared to the coefficient for chr4 inversion cline. **B)** Inversion frequency at sample sites visualized across the black abalone range. **C)** GWAS of latitude against genotype for the 27 largest scaffolds in the reference assembly. Blue and red lines correspond to p-values of 1e-05 (suggested association) and 5e-08 (significant association).

### 2.4 Genetic diversity and demography

Neutral genetic diversity was uniformly high across our sampled abalone. We calculated the average number of differences between pairs of genomes at a site (Θ_π_; _(Tajima, 1989)_ at each discrete site in our study (n=35; Fig 1A). Median Θ_π_ is 0.64 (min: 0.45; max: 0.67), equivalent to ∼ 1 SNP every 150 bp in the genome. While our per-site sample sizes were small (n=3-4), previous work has shown that small sample sizes are most likely to skew estimates of Θ_π_ downward (Subramanian, 2016), resulting in an underestimate of true diversity. We found no clear pattern in the distribution of polymorphism; even the southern sites most impacted by withering syndrome showed genetic diversity on par with the less impacted northern sites (Fig. 5A). In agreement with our observations of high genetic diversity and minimal population structure, we observed that linkage disequilibrium (LD) decays rapidly in these data, indicating extensive recombination and mixing of haplotypes. To explore LD decay, we first defined northern and southern metapopulations each consisting of high coverage (>8X) individuals from northern or southern collection sites in close proximity (Fig. 5B). Within these metapopulations, we observe that the correlation (*r^2^*) between SNPs drops below 0.2, - a common threshold below which variants are not considered strongly ‘linked’ (Hahn, 2018) - within just 50 bp (Fig. 5B).

**Figure 5.**
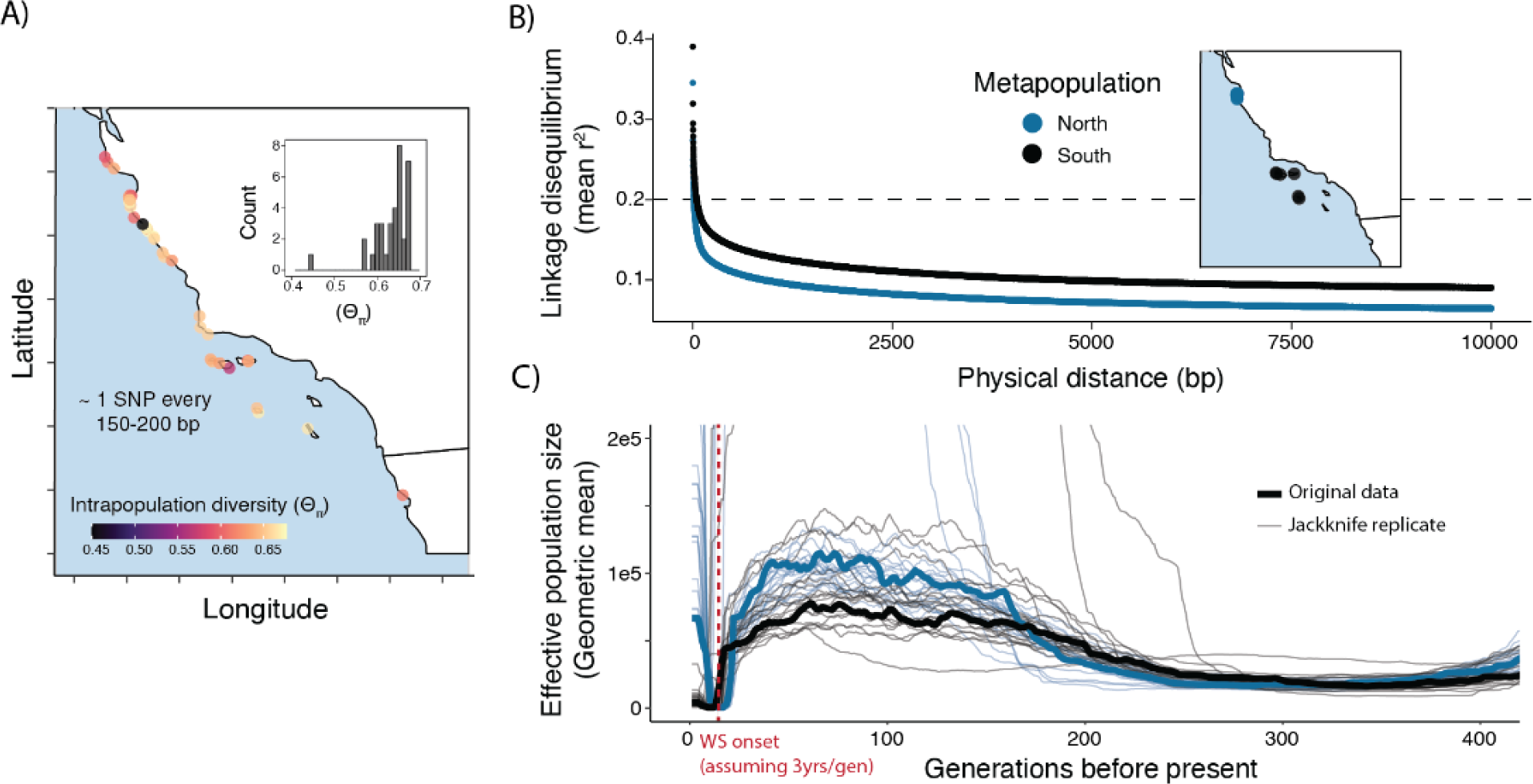
**A)** Intrapopulation genetic diversity (Θ_π_) visualized by location. Inset corresponds to a histogram of the same values plotted as points on the map. **B)** Average linkage disequilibrium plotted by physical distance from a given focal SNP for northern and southern metapopulations **C)** Effective population size through time for metapopulations.

While the results outlined above highlight surprising genetic diversity in today’s populations, genomic inference of demographic history nevertheless shows a severe bottleneck occurring in the species’ recent past. First, by examining coalescent histories with *SMC++* (Terhorst et al., 2017), we identified a decline in both northern and southern metapopulations beginning around 5,000 generations but appearing to stabilize towards the present (Fig. S4). However, As *SMC++* and related methods have difficulty resolving events in the very recent past (<100 generations), we also inferred demographic history with *GONE*, which uses the spectrum of LD between alleles to infer demography in the past 200 generations <u>(Santiago et al. 2020)</u>. With this approach we detect a roughly 99% decline in N_e_ of the southern metapopulation beginning 18 generations ago and reaching a minimum at 10 generations in the past (Fig. 5C). The northern metapopulation had a similar trajectory, with a bottleneck of roughly the same magnitude between 22 generations and 15 generations in the past. (Fig. 5C). For both metapopulations, the *GONE* estimates indicated a recovery beginning within the past five generations.

### 2.5 Prevalence of withering syndrome bacteria and phage hyperparasite

DNA from the bacterial agent of withering syndrome was detected in just 10.5% (n = 14) of sequencing libraries. These libraries spanned our sampling range (Fig. 6B), suggesting that, while not common, WS-RLO is nevertheless sporadically present in black abalone across the species’ range. Indeed, there is no significant association between presence or absence of WS-RLO and latitude (Fig. 6B; Welch two-sample t-test p = 0.90). The phage hyperparasite is similarly scarce, and is observed in just 12.8% (n = 17) of sequencing libraries. The presence or absence of phage shows no significant association with latitude either, suggesting that one is just as likely to observe it in northern sites as southern sites (Fig. 6B; Welch two-sample t-test p = 0.23)

**Figure 6.**
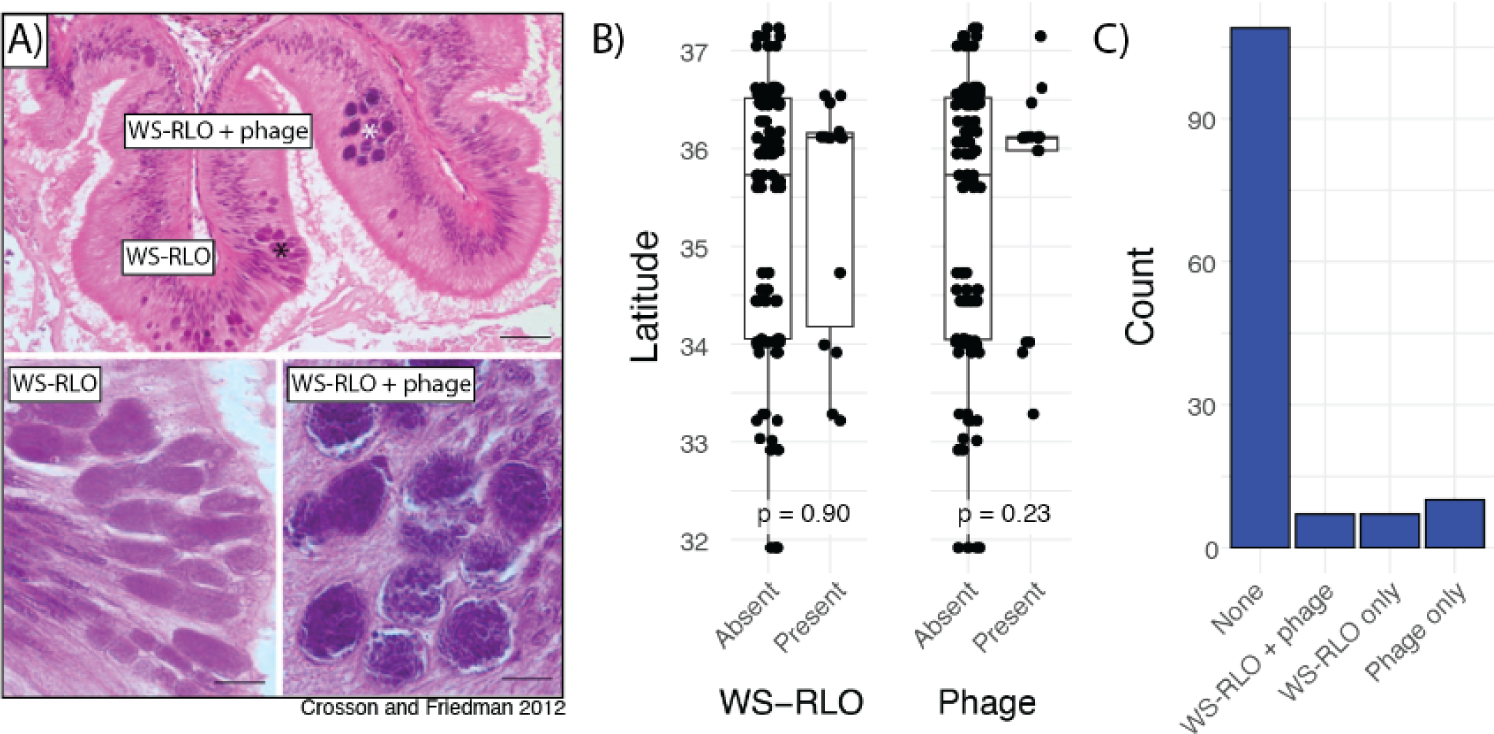
**A)** Representative light micrographs of WS-RLO and phage-infected WS-RLO in the posterior esophagus of black abalone. Figure adapted from (Friedman & Crosson, 2012)**. B)** Association between latitude and presence of WS-RLO or phage. **C)** Observed counts of individuals with for all combinations of WS-RLO and phage presence or absence.

Surprisingly, WS-RLO and phage are not always observed together in the same sample (Fig. 6C). 5.3% of black abalone contain both parasites, while 5.3% and 7.5% contain just WS-RLO or phage, respectively. The absence of WS-RLO in phage positive samples is not eliminated after applying more strict or relaxed detection criteria to both parasites.

## 3 Discussion

Despite experiencing a near-extinction level decline in the recent past, black abalone (*Haliotis cracherodii*) harbor high genetic diversity and exhibit almost no population structure. (Fig. 5A-B). Our estimates of pairwise sequence diversity rank black abalone as more genetically diverse than most vertebrates (Teixeira & Huber, 2021) and more diverse than organisms with similar biology and life history, including western Pacific abalone (Hirase et al., 2021), which maintain high diversity due to their long-term large effective population sizes. This high diversity is also unstructured across their range (Fig 2C; Fig. 3), rejecting previous hypotheses that the negatively buoyant phase and brief larval duration of black abalone would amplify geographic structure (Chambers et al., 2006). Instead, black abalone appear to be panmictic. A similar lack of genetic structure has been documented in wide-ranging broadcast spawning organisms in California, particularly those with high fecundity, extended spawning periods, and intermediate (2-4 weeks) to long (>8 weeks) planktonic larval stages (Dawson, 2001). Therefore, the apparent panmixia of black abalone is consistent with the species’ life history, yet surprising given the catastrophic bottleneck that occurred just decades ago.

While black abalone were seemingly buffered against a significant loss of genetic diversity during their recent decline, their genomes nonetheless retain a signal of a recent, brief, intense population bottleneck that occurred 10-20 generations ago (Fig. 5C). This signal is intriguing, as the bottleneck’s magnitude - a ∼99% decline in both northern and southern metapopulations - agrees with the 99% decline attributed to withering syndrome south of Point Conception, and to a lesser extent, the 0-50% decline in populations north of this boundary (Neuman et al., 2010). While less expected, the signal of decline in the northern metapopulation could be produced by the combination of a true bottleneck effect and a ‘southern’ genomic signature present due to high gene flow across the range (Fig. 3). In addition to magnitude, the timing of the inferred decline roughly aligns with the timeline of WS spread, which was first observed in 1986, approximately twelve generations in the past (Leighton & Boolootian, 1963; Neuman et al., 2010) (Fig. 5C). Our models also indicate recovery in recent generations, which agrees with observations of post-WS population growth at some of the California Channel Islands (Kenner & Yee, 2022) and expansion in northern sites (Miner et al., 2006). While estimates of demographic change in recent history should be interpreted with caution, this concordance with decades of fisheries and academic census data suggest that our genomic analyses accurately reflect the demography of the species, and therefore may be used to guide management strategies.

The only signal of strong genetic clustering in our data set was driven by a 31MB chromosomal inversion present on chr4 (Fig. 2A-C, Fig 4). The inversion is geographically structured, with the derived (low polymorphism) allele increasing in frequency with latitude (Fig. 4B, Fig. S3). Domination of genome-wide structure by a single locus is rare but has been observed in other taxa with high gene flow and segregating chromosomal rearrangements (Luna et al., 2023; Mérot et al., 2021). These inversion polymorphisms may persist in populations if they are evolving under some form of selection. Inversions can link adaptive alleles, for example, and prevent recombination with maladaptive haplotypes, the latter of which is more likely to occur in high gene flow systems (B. Charlesworth & Barton, 2018; Hager et al., 2022; Joron et al., 2011; Kirkpatrick & Barton, 2006). The iversions themselves may also be adaptive, for example if an inversion breakpoint disrupts key genes (Küpper et al., 2016; Villoutreix et al., 2021). While the mechanism by which an inversion influences phenotype will vary, chromosomal inversions are repeatedly implicated in local adaptation, for example through direct phenotype associations (Hager et al., 2022; Nosil et al., 2023; Sanchez-Donoso et al., 2022) or association with environmental variation (Kapun & Flatt, 2019; Mérot et al., 2018; Todesco et al., 2022). While no specific phenotype is associated with the black abalone inversion, the significant increase in inversion frequency in northern latitudes (Fig. 4) and the lack of nucleotide diversity in this northern haplotype (Fig. S3) together suggest evolution under natural selection. Given that the northern populations are more resistant to withering syndrome than populations to the south it is tempting to speculate that the inversion is correlated with this resistance. Latitude, however, is also strongly correlated with other environmental variables, including temperature. Functional data, particularly range-wide transcriptomes and detailed disease phenotypes, will be necessary to determine if the inversion is associated with withering syndrome resistance or an entirely different phenotype.

Although southern black abalone populations have historically been more impacted by withering syndrome, we detected WS-RLO, the agent of withering syndrome, in individuals from both the northern and southern ends of our sampled range (Fig. 6). There was no significant difference in WS-RLO presence according to latitude, indicating that, while uncommon in healthy black abalone, the bacteria has a wide range that includes northern, colder sites (e.g. Carmel, CA) which have not experienced withering syndrome outbreaks (Crosson et al., 2014) (Fig. 6B). The phage hyperparasite, while similarly scarce, also showed no significant association with latitude (Fig. 6B). This raises questions as to whether phage presence attenuates WS severity, as has been suggested previously (Friedman et al., 2014), as the distribution of a disease-attenuating phage may be expected to align with the historic presence of disease. However, complementary measures of phage and pathogen abundance and more thorough disease phenotypes will be necessary to confirm these observations. Additionally, while the lack of spatial pattern in the presence of both parasites is intriguing, accurate identification of WS-RLO and phage is likely constrained by technical barriers. A full WS-RLO reference genome would facilitate more precise detection based on read mapping, and consistent sequencing effort would improve comparisons between samples. Accurately determining the level of endemism of WS-RLO and its association with abiotic (e.g. latitude) and biotic (e.g. phage) factors is key to informing conservation strategy.

Our results provide guidance for the ongoing management of black abalone along the Pacific coast. The high degree of genetic diversity remaining among populations and the sharing of this diversity - and WS-RLO - across the range indicates that translocation of individuals from healthy populations could be a feasible and low-risk recovery plan. However, these findings also set the stage for future work, in particular research into potential adaptive loci (i.e. chr4 inversion) to better design region-specific management strategies and avoid eroding locally adaptive genotypes. Whether or not adaptation is occurring or incipient growth will continue, these data clearly show that substantial genetic variation persists in today’s populations. This finding alone is an encouraging sign for the species’ recovery prospects.

## 4 Methods

### 4.1 Sample collection

We collected samples in a semi-invasive, non-lethal manner by firmly swabbing the foot of the black abalone with a sterile nylon-tipped swab. Individuals were not removed from the substrate, with the intention that no long-term injuries would be inflicted on the animals. In total, we swabbed 150 abalone between January 24^th^ 2020 and February 27^th^ 2022 at 25 sites between Pebble Beach (37.23, -122.42) and Boat House (34.55, -120.61) California (USA), at nine sites on five of the California Channel Islands (San Clemente, Santa Rosa, San Nicolas, Santa Cruz, and San Miguel; USA) and one site in Ensenada, Mexico (Fig. 1; Table S1). Sites in the United States were sampled under NMFS ESA Section 10 Research Permit 18761, while samples from Ensenada, Mexico were provided by a collaborator. At the time of swabbing, none of the abalone sampled displayed external signs of WS. We collected swabs in duplicates and stored them in Longmire buffer (100 mM Tris, 100 mM EDTA, 10 mM NaCl, 0.5% SDS, 0.2% sodium azide) to preserve DNA, and then stored them at 4°C. We recorded the size of each individual, as well as whether it was submerged in water at the time of swabbing, its distance to its nearest neighbor, the number of abalone in its sub-site (i.e. crack or crevice containing one or more individuals), and the primary species found at the sub-site.

### 4.2 DNA extraction

We extracted DNA from one of the two duplicate swabs using a modified version of the DNeasy® Blood and Tissue kit (QIAGEN) optimized to recover DNA from swabs. In brief, we incubated the swabs for 2.5-hours at 56°C in 360 µl of Longmire buffer and 40 µl of 20 mg/mL proteinase K. Following the incubation, we transferred the liquid to a fresh tube, spun the swabs at 13,000 rpm in a centrifuge for 1 min, and then transferred any released liquid to the same tube. We increased the volumes of Buffer AL and ethanol to 400 µl, but the rest of the protocol was unchanged. We eluted the DNA in 50 µl of Buffer EB (10 mM Tris), and used 1 µl of extract to quantify DNA concentration with the Qubit dsDNA HS Assay Kit (Invitrogen). We repeated the DNA extraction using the duplicate swab if an extractions DNA concentration was too low for quantification. We stored the extractions at -20°C.

### 4.3 Library preparation and sequencing

We prepared the DNA extracts into sequencing libraries following the NEBNext Ultra II FS DNA Library Prep Kit (NEB) according to manufacturers’ instructions but replaced the NEBNext Adapters with Y-Adapters. We incubated the samples for five minutes during the enzymatic fragmentation step, performed a double-sided size selection (first clean at 0.26 X and second clean at 0.11 X), and amplified the libraries for 6-8 using dual unique indexes. We eluted each library in 21 µl of 0.1 X TE and quantified the DNA concentration using the Qubit dsDNA HS Assay Kit (Invitrogen) and fragment length on a Fragment Analyzer (Agilent). We then screened each library via low coverage sequencing on an Illumina NextSeq 550 (2 x 150 bp). Libraries were then sequenced on an Illumina Novaseq 6000 (2 x 150 bp) with a target depth of 10-20X genome wide coverage.

### 4.4 Mapping and variant calling

We generated a composite reference genome in order to accurately map off-target reads such as those contributed by WS bacteria and phage hyperparasite present in our DNA swabs. This composite reference genome included the black abalone reference (Orland et al., 2022) (GenBank accession number GCA_022045235.1), the bacterial *Candidatus Xenohaliotis californiensis* 16S ribosomal RNA gene (also referred to as the withering syndrome rickettsia-like organism or WS-RLO; GenBank accession number AF133090.2) and the WS-RLO phage genome (also referred to as the RLO variant or RLOv; GenBank accession number KY296501.1).

We used the *snpArcher* workflow (Mirchandani et al., 2024), an accelerated workflow for variant calling, to generate high quality variant calls for downstream analysis. Briefly, sequencing reads were first trimmed of adapter sequence using *fastp* (Chen et al., 2018) and aligned to the composite genome using *bwa mem (Li & Durbin, 2009)*. We called individual variants with Sentieon (Kendig et al., 2019) Haplotyper and we performed joint genotyping using Sentieon Genotyper to produce a multisample VCF (variant call format) file. Using *bcftools (Danecek et al., 2021)*, we removed individuals with <2x sequencing depth, sites with minor allele frequency < 0.01, and sites with > 75% missing data. We additionally removed sites that failed a set of standard GATK hard filtering thresholds (Van der Auwera et al., 2013), defined in the *snpArcher* workflow. We additionally removed all indels and retained only biallelic SNPs for downstream analysis, resulting in 66,776,934 SNPs.

### 4.5 Population structure analyses

To explore population structure via principal components analysis (PCA), we selected unlinked (>1000 bp apart) biallelic SNPs with minor allele frequency > 0.025 followed by random subsampling to speed up computation, resulting in 998,975 total variants. Following this we calculated PCAs at the whole genome level and at genomic regions of interest using the function *allel.pca()* from *scikit-allel v.1.3.6 (Miles, 2015)*. To further explore the contributions of particular genomic regions to PCA clustering in an unbiased fashion, we conducted a local principal components analysis across the genome via *lostruct (Li & Ralph, 2019)*. Prior to running this analysis, we refined our full variant dataset to retain only SNPs with minor allele frequency > 0.05 and removed sites with >50% missing data, resulting in 25,036,332 SNPs. Finally, we ran *lostruct*, calculating principal component analyses in 5kbp windows.

### 4.6 Chromosomal inversion detection

Based on the results of our preliminary PCA, we scanned our reference genome assembly for potential errors or structural anomalies that might be driving the observed signal. We generated a chromatin-interaction map (contact map) using the Omni-C sequencing data generated for the genome assembly. We inspected the contact map visually and identified two regions in chr4 (scaffold 4) with a high intensity signal off-diagonal that resembled an inversion (Fig. S1). To confirm that this region was an inversion, we searched the PacBio HiFi data generated for the genome assembly for structural variants using *Sniffles* with default parameters (Sedlazeck et al., 2018). We then calculated linkage disequilibrium (LD) for all pairs of SNPs on chr4 using *plink v1.90b7 --r2 inter-chr gz dprime yes-really --ld-window-r2 0* (Purcell et al., 2007).

We tested for associations between the identified chromosomal inversion and environmental variables in a GWAS framework using *EMMAX (Kang et al., 2010)*. We thinned the genotype data to one biallelic SNP every 1000 bp in order to speed up runtimes while still sampling frequently along the genome. We also generated a kinship matrix for our samples based on an inversion-masked genome using *plink2 –make-king square (Chang et al., 2015)*. Finally, to run *EMMAX*, we supplied the genotype data, the kinship matrix, and the first two genetic principal components from the no-inversion PCA as predictors, and supplied a response variable (e.g. temperature) for association testing.

### 4.7 Isolation-by-distance (IBD) and Isolation-by-environment (IBE)

To assess relationships between genetic and ecological divergence, we obtained several physical and environmental variables for each site. We calculated physical distance between sites using the function *distm(…,fun = distVincentyEllipsoid)* from the R package *geosphere* (Hijmans, 2022). We modeled connectivity between sites using mathematical particles in a Regional Oceanic Modeling System (ROMS). Projections were based on the latitude and longitude where particles were released (i.e. donor site) and arrived (i.e. settlement site) after planktonic larval durations (PLD) of 5, 10, and 15 days.

We collected annual average sea surface temperatures (SST) from loggers at each site from as early as 1999 to 2021. Temperature loggers (HOBO TidbiT and HOBO Pendant from Onset Computer Corporation) were deployed in the low intertidal zone and recorded temperature every 15 minutes. We either downloaded loggers in the field using Onset app (HOBOconnect) or collected and downloaded loggers post-field using Onset software (HOBOware), and exported these data as an ASCII file. We processed the data to separate air temperature and water temperature by comparing the data with tide charts and removing temperatures at times where the tidal height was predicted to be lower than the tidal location of the logger. Daily mean sea water temperature was calculated and then averaged to annual mean sea water temperature.

We also obtained monthly air temperature (AT) values using the function *worldclim_country(“USA”, var=“tavg”, path=tempdir(),res=5)* from the R package *geodata* (Hijmans et al., 2022), selected regions corresponding to our sample locations with *terra::extract(…,method = ‘bilinear’)*, and calculated the mean annual temperature for each site. pH data were averaged annually from monthly collections at a 3 × 3 km resolution from 1990 to 2010 but should reflect the average variation pH among sites (Cheresh & Fiechter, 2020). In order to transform SST, AT, pH per-site values into differences between sites, we generated distance matrices with the base R function *dist(…, method=’manhattan’)*. Following this, we searched for associations between these aspects of environmental distance using the base R function *prcomp*. After observing strong loadings of all variables on PC1 (Fig S2), we thereafter summarized environmental dissimilarity by a site pair’s PC1 score.

Finally, we calculated genetic distance between sites in 10 kb windows using Hudson’s F_ST_ via the *scikit-allel v.1.3.6* function *allel.windowed_hudson_fst()*.

### 4.8 Genetic diversity and demographic inference

We quantified genetic diversity (Θ_ⲡ_) in 1kb windows using the function *allel.windowed_diversity()* from *scikit-allel v.1.3.6*. To obtain summaries of linkage disequilibrium we ran *PopLDdecay* (Zhang et al., 2019) on whole genome data with the exception of the chr4 inversion.

We used *GONE*, an LD-spectrum based demographic inference tool, to infer population size through recent time (Santiago et al., 2020). In order to gain sufficient resolution and confidence in our LD spectrum, we designated two metapopulations, ‘north’ and ‘south’, consisting of 18 and 12 samples respectively with sequencing coverage greater than 8, existing in close proximity, and exhibiting minimal genetic structure. To reduce runtime we thinned our SNP dataset to ∼1 random variant every 1 kb, while masking the chr4 inversion. Finally, we converted this input data to the *plink* ped/map format with *plink*. We ran *GONE* on this dataset with *script_GONE.sh*, setting default parameters with the exception of *NGEN=2000 NBIN=400 maxNSNP=50000 hc=0.10*. For lack of fine-scale recombination rate data in this species, we assumed a constant recombination rate of 1 cM/Mb with the parameter *cMMb = 1*. We re-ran *GONE* on genomic subsamples consisting of randomly sampled chromosomes, ranging from 50%-80% of the genome.

We also used *SMC++* (Terhorst et al., 2017) to infer longer term changes in population size. For this analysis we used the same northern and southern metapopulations, but because *SMC++* relies on an accurate identification of heterozygous sites as well as the site frequency spectrum (SFS), we generated conservative ‘mappability’ and ‘depth’ masks for the input data. To generate a mask of low-mappability regions, we used *GenMap (Pockrandt et al., 2020)* on the reference assembly with the parameters ‘*-K 150 -E 4’*, and retained low-quality regions where kmers mapped to three or more places in the genome. We then used a custom script (*bamdepth2bed.py*) to generate depth masks for each metapopulation to indicate where more than 30% of samples had 5 or less reads mapping to a position, and therefore where genotype information might be unreliable. We then converted the vcf files to *SMC++* format using *vcf2smc*, designated the highest coverage sample of each metapopulation as the ‘Distinguished Individual’ (DI) while masking the inversion, low mappability regions, and low depth regions. Finally we ran *smc++ estimate --timepoints 1e3 1e6 --knots 7 --spline piecewise* to generate demographic histories, providing the human germline mutation rate of 2.5e-8 (Lindsay et al., 2019) for lack of any suitable mollusk rate. We then generated bootstrap resampled datasets using a custom script (‘*SMC_bootstrap_BW.py*’) and reran the above pipeline for each replicate.

### 4.9 Associations with withering syndrome bacteria and phage hyperparasite

We determined the presence of WS-RLO in our DNA swab libraries using the eDNA community profiling tool *tronko (Pipes & Nielsen, 2022)*. Specifically, for each library we selected all paired-end reads not mapping to the *H. cracherodii* genome. We then provided the sequence reads and the pre-built 16S DNA reference database (https://zenodo.org/records/7407318) to *tronko* specifying a least common ancestor (LCA) cutoff of 4 (*-c 4*). After examining all hits to taxa in the order *Rickettsiales*, which are obligatory intracellular parasites, we then designated reads mapping to *Candidatus Xenohaliotis californicus 16S* (AF133090.2) and *Haplosporidium sp. endosymbiont AbFoot 16S* (AJ319724.1) as true “positives” for WS-RLO presence in our samples.

To detect the phage hyperparasite of WS-RLO, we capitalized on the availability of a high quality reference genome (KY296501.1). We identified reads mapping to the 35,728 bp genome (see 2.4), and designated libraries with >25% of the genome covered by one or more reads as “positive” for the phage, assuming such a profile is unlikely to be generated by spurious mapping from other taxa present in our swabs. This heuristic also allowed us to identify libraries that had disproportionate mapping along the phage genome despite lower sequencing effort, avoiding potential false negatives.

## Author contributions

Chloe Orland, Beth Shapiro, and Peter Raimondi conceptualized the study. Joshua Kapp, Chloe Orland, Nathaniel Fletcher and Karah Ammann coordinated and completed the sampling, and laboratory work was done by Chloe Orland and Joshua Kapp. ROMS data were provided by Peter Raimondi. Bioinformatic and data analyses were conducted by Brock Wooldridge, Chloe Orland, Erik Enbody, Merly Escalona, and Cade Mirchandani. Brock Wooldridge and Chloe Orland wrote the manuscript with input and approval from all authors.

## Acknowledgments

The authors thank the field sampling teams: Laura Anderson, Maya George, Christy Bell, Melissa Douglas, Rani Gaddam, Mia Cortez, Lexi Necarsulmer, Dan Richards, Frankie Gerraty, Kim Parisi, Rhys Evans, Steve Whitaker, Torrey Gorra, Lindsay Cullen, Josh Sprague, Brian Hong, Dominic Richards, and Jessica Bredvik. They also thank Hailey Nava for her help processing samples and the UC Santa Cruz Paleogenomics Laboratory for their support in the lab. Finally, we are grateful to Richard E. Green for his valuable advice, to Alicia Abadia Cordoso and Rodrigo Beas for providing samples from Mexico, and to Jerome Fiechter for modeling environmental variables for our collection sites. This research was supported by a California Conservation Genomics Project grant. Erik Enbody and Russell Corbett-Detig were supported by the National Institutes of Health (Grant No. R35GM128932). Brock Wooldridge was supported by the National Science Foundation Ocean Sciences Postdoctoral Fellowship (No. 2307479).

## Data Accessibility and Benefit Sharing Statement

All sequence data are available on the NCBI Short Read Archive (SRA) database under BioProject PRJNA982170.

Environmental metadata generated or synthesized for this study, including water temperature, pH, and ROMS modeled connectivity, are available on the Dryad database at 10.5061/dryad.r7sqv9skq.

## Supplement Figures

**Figure S1.**
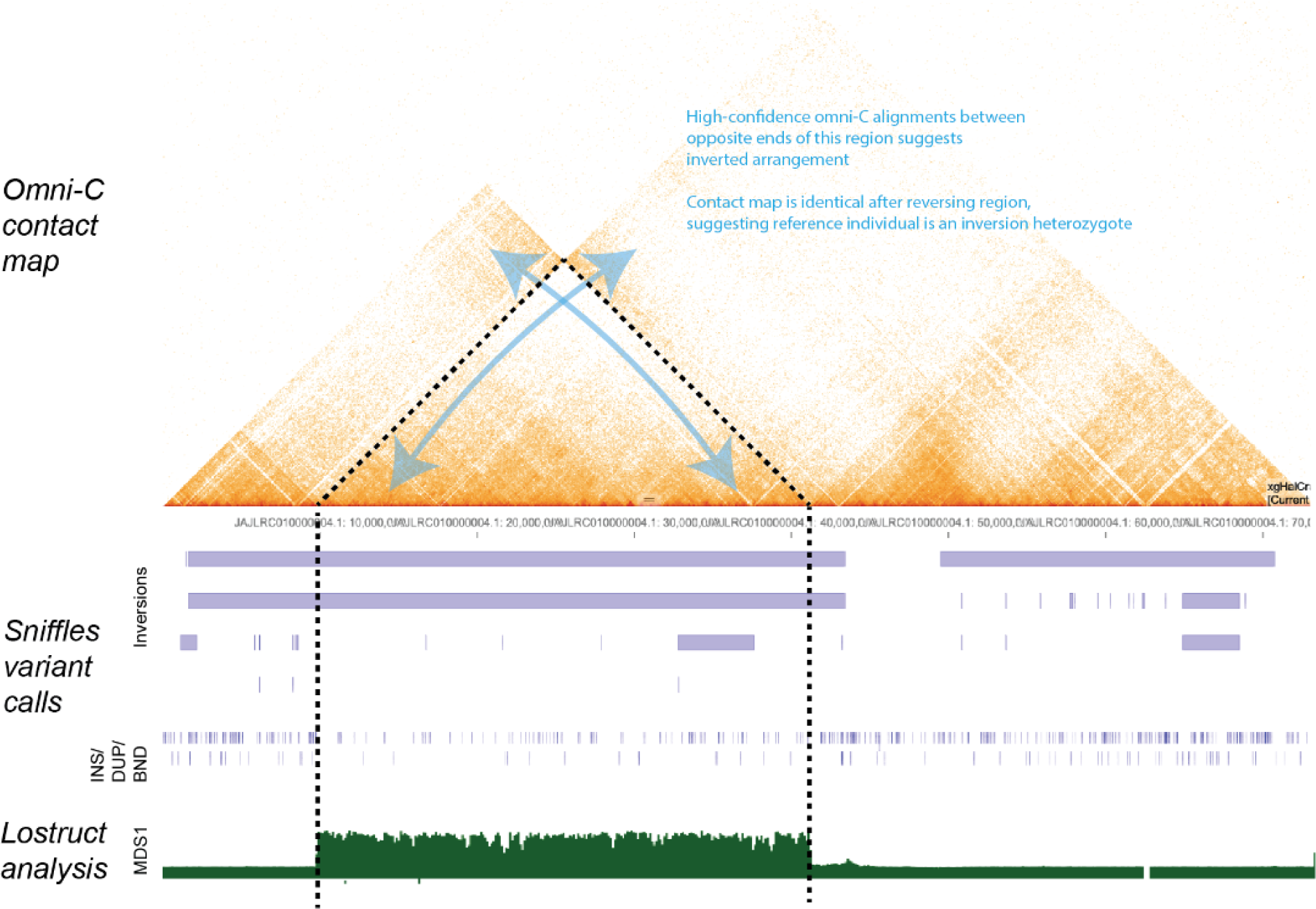
Evidence for a chromosomal inversion in chr4. Top panel displays Omni-C linkage mapping data generated for the reference assembly (Orland et al., 2022) showing an unsolvable region in chr4 that suggests the preference of a chromosomal inversion. Middle panel shows structural variants genotyped based on PacBio long-read variant calls with *Sniffles (Sedlazeck et al., 2018*). INS/DUP/BND refers to insertions, duplications, and translocations, respectively. Proposed inversions overlap imperfectly with boundaries suggested by Omni-C, indicating some uncertainty with inversion coordinates. Bottom panel displays *lostruct* results for chr4 (Li and Ralph, 2019). Each point corresponds to the local genomic structure in 5000 bp genomic windows, with the “MDS1” position roughly representing deviation in the window from genome-wide structure.

**Figure S2.**
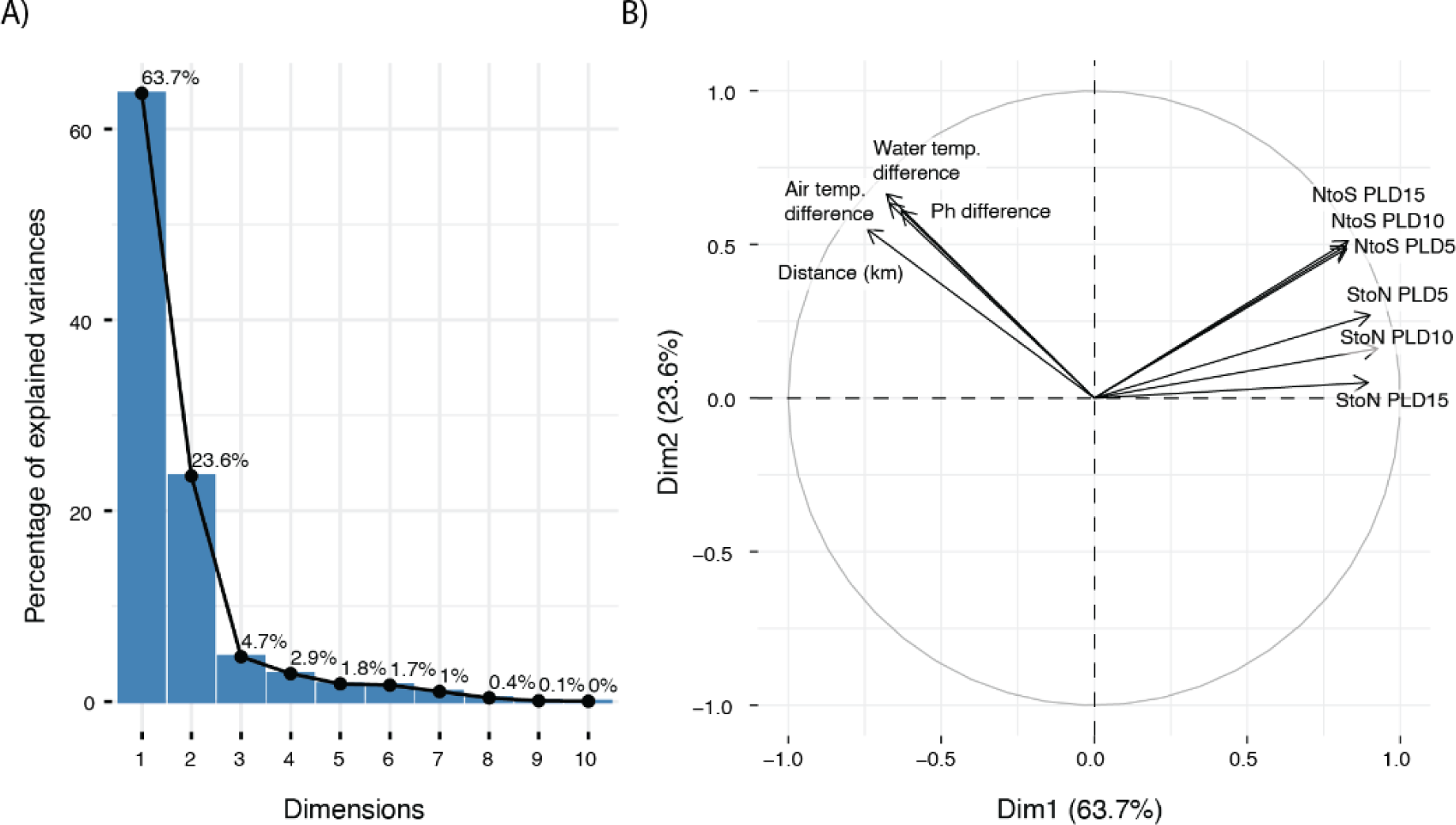
Results of PCA on environmental and physical variables associated with each site. A) Scree plot indicating proportion of variance explained by the first 10 PCs. B) Variable loadings indicating the associations between each variable and the first two PCs.

**Figure S3.**
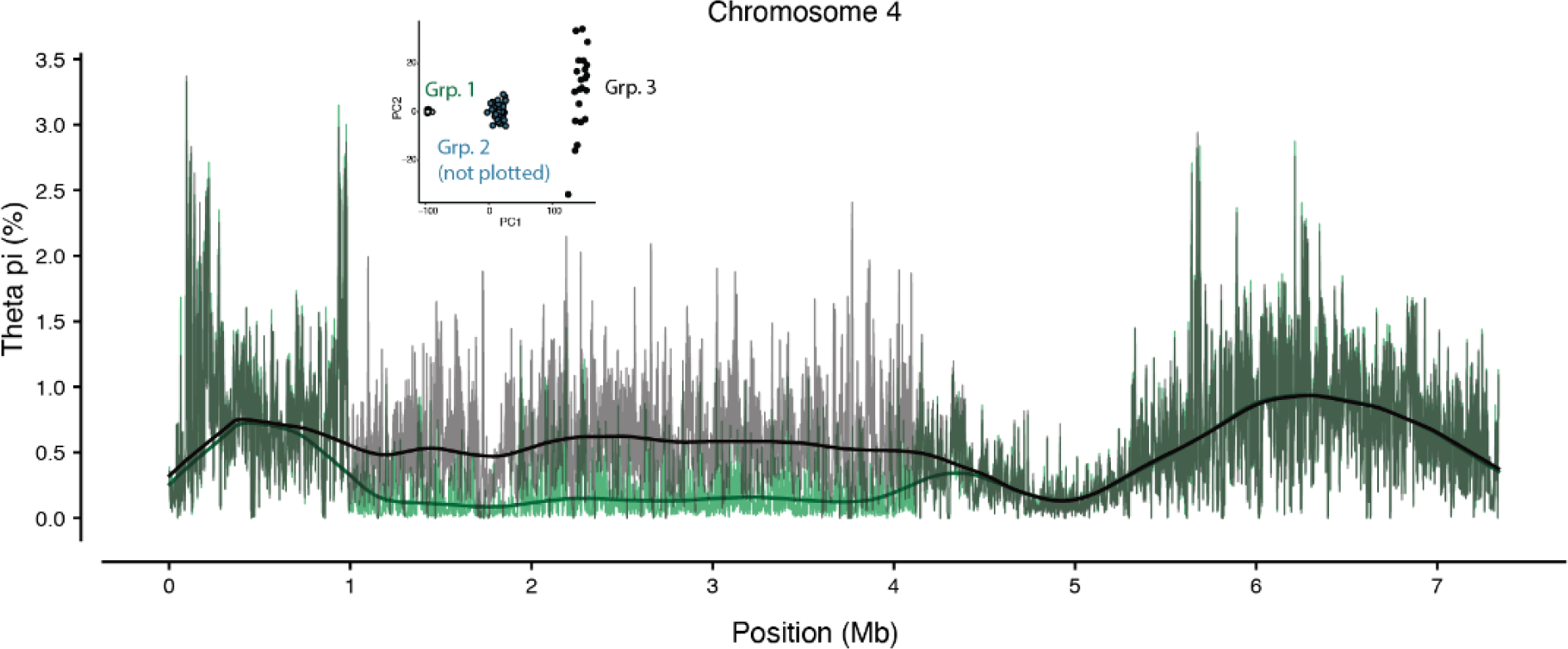
Genetic diversity within inversion haplotypes. Diversity (Θ_π_) is plotted in 10kb windows, with bold lines representing the loess-smoothed fit. The inset reproduces the PCA seen in Fig. 2B to indicate which inversion clusters are being plotted here.

**Figure S4.**
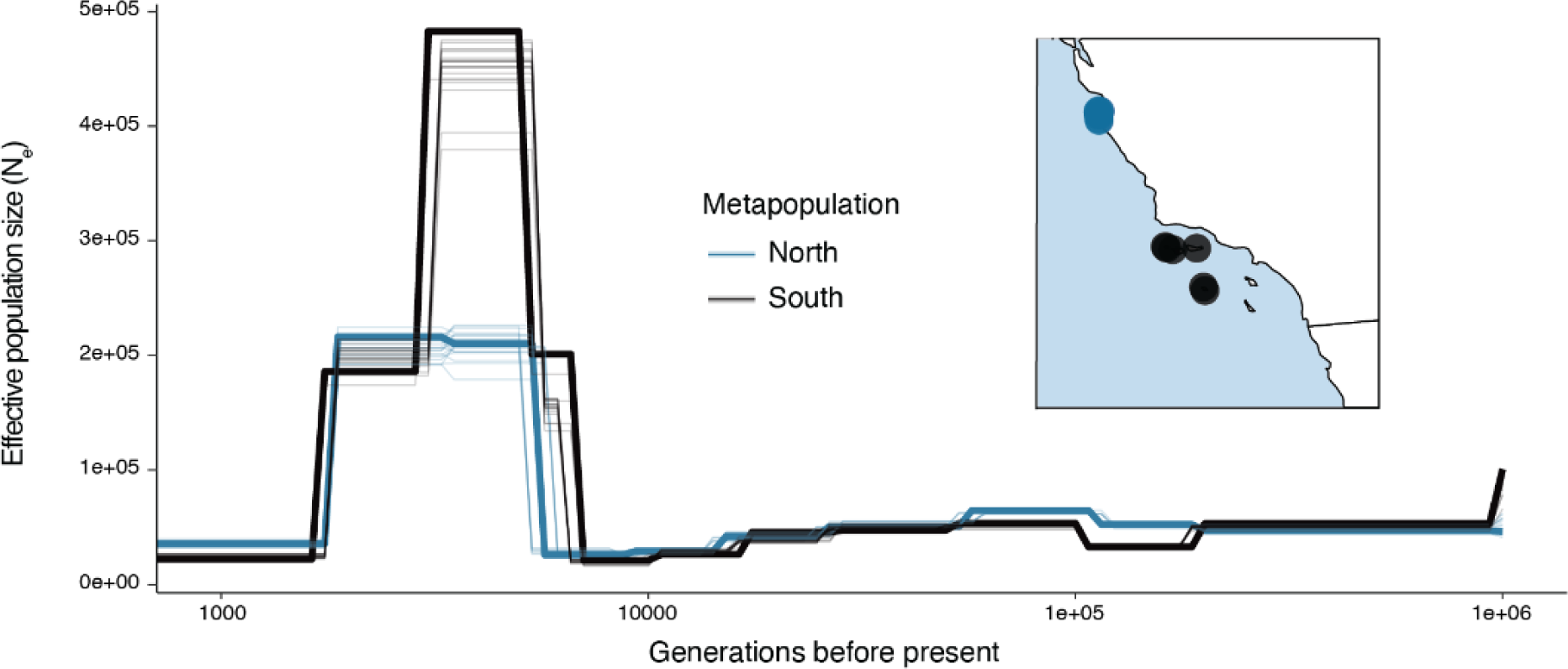
Effective population size through time for northern and southern metapopulations. Analysis performed with SMC++ (Terhorst et al., 2016) and assuming a germline mutation rate of 2.5e-8.

## Tables

**Extended Data Table S1.** List of swabbed samples and collection locations. <u>Alternate name</u> = more informative shorthand name, where applicable; <u>Site</u> = collection site shorthand; <u>Longitude,</u> <u>Latitude</u> = coordinates with ∼1km accuracy; <u>QCpassing</u> = whether or not sample generated sufficient endogenous DNA for genomic analyses.

## References

Agrawal, A. F., & Whitlock, M. C. (2012). Mutation load: the fitness of individuals in populations where deleterious alleles are abundant. Annual Review of Ecology, Evolution, and Systematics, 43(1), 115–135.

Altstatt, J. M., Ambrose, R. F., Engle, J. M., Haaker, P. L., Lafferty, K. D., & Raimondi, P. T. (1996). Recent declines of black abalone *Haliotis cracherodii* on the mainland coast of central California. Marine Ecology Progress Series, 142, 185–192.

Ben-Horin, T., Lenihan, H. S., & Lafferty, K. D. (2013). Variable intertidal temperature explains why disease endangers black abalone. Ecology, 94(1), 161–168.

Bentz, L., & Braje, T. J. (2017). Sea of prosperity: foundations of the California commercial abalone fishery. International Journal of Historical Archaeology, 21(3), 598–622.

Braje, T. J., Costello, J. G., Erlandson, J. M., & DeLong, R. (2014). Of seals, sea lions, and abalone: the archaeology of an historical multiethnic base camp on San Miguel Island, California. Historical Archaeology, 48(2), 122–142.

Brokordt, K., González, R., Farías, W., Winkler, F. E., & Lohrmann, K. B. (2017). First insight into the heritable variation of the resistance to infection with the bacteria causing the withering syndrome disease in *Haliotis rufescens* abalone. Journal of Invertebrate Pathology, 150, 15–20.

Chambers, M. D., VanBlaricom, G. R., Hauser, L., Utter, F., & Friedman, C. S. (2006). Genetic structure of black abalone (*Haliotis cracherodii*) populations in the California islands and central California coast: Impacts of larval dispersal and decimation from withering syndrome. Journal of Experimental Marine Biology and Ecology, 331(2), 173–185.

Chang, C. C., Chow, C. C., Tellier, L. C., Vattikuti, S., Purcell, S. M., & Lee, J. J. (2015). Second-generation PLINK: rising to the challenge of larger and richer datasets. GigaScience, 4, 7.

Charlesworth, B. (2009). Fundamental concepts in genetics: effective population size and patterns of molecular evolution and variation. Nature Reviews. Genetics, 10(3), 195–205.

Charlesworth, B., & Barton, N. H. (2018). The spread of an inversion with migration and selection. Genetics, 208(1), 377–382.

Charlesworth, D., & Willis, J. H. (2009). The genetics of inbreeding depression. Nature Reviews. Genetics, 10(11), 783–796.

Chen, S., Zhou, Y., Chen, Y., & Gu, J. (2018). fastp: an ultra-fast all-in-one FASTQ preprocessor. Bioinformatics, 34(17), i884–i890.

Cheresh, J., & Fiechter, J. (2020). Physical and biogeochemical drivers of alongshore pH and oxygen variability in the California current system. Geophysical Research Letters, 47(19). 10.1029/2020gl089553

Cox, K. W. (1962). California abalones, family Haliotidae. Fishery Bulletin, 118.

Crosson, L. M., & Friedman, C. S. (2018). Withering syndrome susceptibility of northeastern Pacific abalones: A complex relationship with phylogeny and thermal experience. Journal of Invertebrate Pathology, 151, 91–101.

Crosson, L. M., Wight, N., VanBlaricom, G. R., Kiryu, I., Moore, J. D., & Friedman, C. S. (2014). Abalone withering syndrome: distribution, impacts, current diagnostic methods and new findings. Diseases of Aquatic Organisms, 108(3), 261–270.

Danecek, P., Bonfield, J. K., Liddle, J., Marshall, J., Ohan, V., Pollard, M. O., Whitwham, A., Keane, T., McCarthy, S. A., Davies, R. M., & Li, H. (2021). Twelve years of SAMtools and BCFtools. GigaScience, 10(2). 10.1093/gigascience/giab008

Dawson, M. N. (2001). Phylogeography in coastal marine animals: a solution from California? Journal of Biogeography, 28(6), 723–736.

Erlandson, J. M., Rick, T. C., Braje, T. J., Steinberg, A., & Vellanoweth, R. L. (2008). Human impacts on ancient shellfish: a 10,000 year record from San Miguel Island, California. Journal of Archaeological Science, 35(8), 2144–2152.

Frankham, R., Lees, K., Montgomery, M. E., England, P. R., Lowe, E. H., & Briscoe, D. A. (1999). Do population size bottlenecks reduce evolutionary potential? Animal Conservation, 2(4), 255–260.

Friedman, C. S., Andree, K. B., Beauchamp, K. A., Moore, J. D., Robbins, T. T., Shields, J. D., & Hedrick, R. P. (2000). “*Candidatus* Xenohaliotis californiensis”, a newly described pathogen of abalone, Haliotis spp., along the west coast of North America. International Journal of Systematic and Evolutionary Microbiology, 50, 847–855.

Friedman, C. S., & Crosson, L. M. (2012). Putative phage hyperparasite in the rickettsial pathogen of abalone, “Candidatus Xenohaliotis californiensis.” Microbial Ecology, 64(4), 1064–1072.

Friedman, C. S., Wight, N., Crosson, L. M., Vanblaricom, G. R., & Lafferty, K. D. (2014). Reduced disease in black abalone following mass mortality: phage therapy and natural selection. Frontiers in Microbiology, 5, 78.

Grossen, C., Guillaume, F., Keller, L. F., & Croll, D. (2020). Purging of highly deleterious mutations through severe bottlenecks in Alpine ibex. Nature Communications, 11(1), 1001.

Gruenthal, K. M., & Burton, R. S. (2008). Genetic structure of natural populations of the California black abalone (*Haliotis cracherodii* Leach, 1814), a candidate for endangered species status. Journal of Experimental Marine Biology and Ecology, 355(1), 47–58.

Haas, H., Braje, T. J., Edwards, M. S., Erlandson, J. M., & Whitaker, S. G. (2019). Black abalone (*Haliotis cracherodii*) population structure shifts through deep time: Management implications for southern California’s northern Channel Islands. Ecology and Evolution, 9(8), 4720–4732.

Hager, E. R., Harringmeyer, O. S., Wooldridge, T. B., Theingi, S., Gable, J. T., McFadden, S., Neugeboren, B., Turner, K. M., Jensen, J. D., & Hoekstra, H. E. (2022). A chromosomal inversion contributes to divergence in multiple traits between deer mouse ecotypes. Science, 377(6604), 399–405.

Hahn, M. W. (2018). Molecular Population Genetics. Oxford University Press.

Hamm, D. E., & Burton, R. S. (2000). Population genetics of black abalone, *Haliotis cracherodii*, along the central California coast. Journal of Experimental Marine Biology and Ecology, 254(2), 235–247.

Hijmans, R. (2022). *gepsphere: Spherical Trigonometry* (Version R package version 1.5-18). https://CRAN.R-project.org/package=geosphere

Hirase, S., Yamasaki, Y. Y., Sekino, M., Nishisako, M., Ikeda, M., Hara, M., Merilä, J., & Kikuchi, K. (2021). Genomic evidence for speciation with gene flow in broadcast spawning marine invertebrates. Molecular Biology and Evolution, 38(11), 4683–4699.

Hoffmann, A. A., Sgrò, C. M., & Kristensen, T. N. (2017). Revisiting adaptive potential, population size, and conservation. Trends in Ecology & Evolution, 32(7), 506–517.

Joron, M., Frezal, L., Jones, R. T., Chamberlain, N. L., Lee, S. F., Haag, C. R., Whibley, A., Becuwe, M., Baxter, S. W., Ferguson, L., Wilkinson, P. A., Salazar, C., Davidson, C., Clark, R., Quail, M. A., Beasley, H., Glithero, R., Lloyd, C., Sims, S., … ffrench-Constant, R. H. (2011). Chromosomal rearrangements maintain a polymorphic supergene controlling butterfly mimicry. Nature, 477(7363), 203–206.

Kang, H. M., Sul, J. H., Service, S. K., Zaitlen, N. A., Kong, S.-Y., Freimer, N. B., Sabatti, C., & Eskin, E. (2010). Variance component model to account for sample structure in genome-wide association studies. Nature Genetics, 42(4), 348–354.

Kapun, M., & Flatt, T. (2019). The adaptive significance of chromosomal inversion polymorphisms in *Drosophila melanogaster*. Molecular Ecology, 28(6), 1263–1282.

Kardos, M., Armstrong, E. E., Fitzpatrick, S. W., Hauser, S., Hedrick, P. W., Miller, J. M., Tallmon, D. A., & Funk, W. C. (2021). The crucial role of genome-wide genetic variation in conservation. Proceedings of the National Academy of Sciences of the United States of America, 118(48). 10.1073/pnas.2104642118

Karpov, K. A., Haaker, P. L., Taniguchi, I. K., & Rogers-Bennett, L. (2000). Serial depletion and the collapse of the California abalone (Haliotis spp.) fishery. Can. J. Fish. Aquat. Sci., 130, 11–24.

Keller, L. F., & Waller, D. M. (2002). Inbreeding effects in wild populations. Trends in Ecology & Evolution, 17(5), 230–241.

Kelley, K., & Francis, H. (2003). Abalone shell Buffalo People: Navajo narrated routes and pre-Columbian archaeological sites. New Mexico Historical Review, 78(1), 3.

Kendig, K. I., Baheti, S., Bockol, M. A., Drucker, T. M., Hart, S. N., Heldenbrand, J. R., Hernaez, M., Hudson, M. E., Kalmbach, M. T., Klee, E. W., Mattson, N. R., Ross, C. A., Taschuk, M., Wieben, E. D., Wiepert, M., Wildman, D. E., & Mainzer, L. S. (2019). Sentieon DNASeq variant calling workflow demonstrates strong computational performance and accuracy. Frontiers in Genetics, 10, 736.

Kenner, M. C., & Yee, J. L. (2022). *Black Abalone surveys at Naval Base Ventura County, San Nicolas Island, California—2021, annual report* (2022-1107). U.S. Geological Survey. 10.3133/ofr20221107

Kirkpatrick, M., & Barton, N. (2006). Chromosome inversions, local adaptation and speciation. Genetics, 173(1), 419–434.

Küpper, C., Stocks, M., Risse, J. E., Dos Remedios, N., Farrell, L. L., McRae, S. B., Morgan, T. C., Karlionova, N., Pinchuk, P., Verkuil, Y. I., Kitaysky, A. S., Wingfield, J. C., Piersma, T., Zeng, K., Slate, J., Blaxter, M., Lank, D. B., & Burke, T. (2016). A supergene determines highly divergent male reproductive morphs in the ruff. Nature Genetics, 48(1), 79–83.

Lafferty, K. D., & Kuris, A. M. (1993). Mass mortality of abalone *Haliotis cracherodii* on the California Channel Islands: tests of epidemiological hypotheses. Mar Ecol Prog Ser, 96, 239–248.

Leighton, D., & Boolootian, R. A. (1963). Diet and growth in the black abalone, *Haliotis cracherodii*. Ecology, 44(2), 228–238.

Li, H., & Durbin, R. (2009). Fast and accurate short read alignment with Burrows-Wheeler transform. Bioinformatics, 25(14), 1754–1760.

Li, H., & Ralph, P. (2019). Local PCA Shows How the Effect of Population Structure Differs Along the Genome. Genetics, 211(1), 289–304.

Lindsay, S. J., Rahbari, R., Kaplanis, J., Keane, T., & Hurles, M. E. (2019). Similarities and differences in patterns of germline mutation between mice and humans. Nature Communications, 10(1), 4053.

Luna, L. W., Williams, L. M., Duren, K., Tyl, R., Toews, D. P. L., & Avery, J. D. (2023). Whole genome assessment of a declining game bird reveals cryptic genetic structure and insights for population management. Molecular Ecology, 32(20), 5498–5513.

Lynch, M., Conery, J., & Burger, R. (1995). Mutation accumulation and the extinction of small populations. The American Naturalist, 146(4), 489–518.

Mérot, C., Berdan, E. L., Babin, C., Normandeau, E., Wellenreuther, M., & Bernatchez, L. (2018). Intercontinental karyotype–environment parallelism supports a role for a chromosomal inversion in local adaptation in a seaweed fly. Proceedings of the Royal Society B: Biological Sciences, 285(1881), 20180519.

Mérot, C., Berdan, E. L., Cayuela, H., Djambazian, H., Ferchaud, A.-L., Laporte, M., Normandeau, E., Ragoussis, J., Wellenreuther, M., & Bernatchez, L. (2021). Locally adaptive inversions modulate genetic variation at different geographic scales in a seaweed fly. Molecular Biology and Evolution, 38(9), 3953–3971.

Miles, A. (2015). *scikit-allel: A Python package for exploring and analysing genetic variation data* (Version 1.3.6). Github. https://github.com/cggh/scikit-allel

Miner, C. M., Altstatt, J. M., Raimondi, P. T., & Minchinton, T. E. (2006). Recruitment failure and shifts in community structure following mass mortality limit recovery prospects of black abalone. Marine Ecology Progress Series, 327, 107–117.

Mirchandani, C. D., Shultz, A. J., Thomas, G. W. C., Smith, S. J., Baylis, M., Arnold, B., Corbett-Detig, R., Enbody, E., & Sackton, T. B. (2024). A fast, reproducible, high-throughput variant calling workflow for population genomics. Molecular Biology and Evolution, 41(1). 10.1093/molbev/msad270

Morse, D. E., Hooker, N., Duncan, H., & Jensen, L. (1979). Ggr-aminobutyric acid, a neurotransmitter, induces planktonic abalone larvae to settle and begin metamorphosis. Science, 204(4391), 407–410.

Neuman, M., Tissot, B., & VanBlaricom, G. (2010). Overall status and threats assessment of black abalone (*Haliotis Cracherodii* Leach, 1814) populations in California. Journal of Shellfish Research, 29(3), 577–586.

Nosil, P., Soria-Carrasco, V., Villoutreix, R., De-la-Mora, M., de Carvalho, C. F., Parchman, T., Feder, J. L., & Gompert, Z. (2023). Complex evolutionary processes maintain an ancient chromosomal inversion. Proceedings of the National Academy of Sciences of the United States of America, 120(25), e2300673120.

Orland, C., Escalona, M., Sahasrabudhe, R., Marimuthu, M. P. A., Nguyen, O., Beraut, E., Marshman, B., Moore, J., Raimondi, P., & Shapiro, B. (2022). A draft reference genome assembly of the critically endangered black abalone, *Haliotis cracherodii*. The Journal of Heredity, 113(6), 665–672.

Pipes, L., & Nielsen, R. (2022). A rapid phylogeny-based method for accurate community profiling of large-scale metabarcoding datasets. In bioRxiv (p. 2022.12.06.519402). 10.1101/2022.12.06.519402

Pockrandt, C., Alzamel, M., Iliopoulos, C. S., & Reinert, K. (2020). GenMap: ultra-fast computation of genome mappability. Bioinformatics, 36(12), 3687–3692.

Purcell, S., Neale, B., Todd-Brown, K., Thomas, L., Ferreira, M. A. R., Bender, D., Maller, J., Sklar, P., de Bakker, P. I. W., Daly, M. J., & Sham, P. C. (2007). PLINK: a tool set for whole-genome association and population-based linkage analyses. American Journal of Human Genetics, 81(3), 559–575.

Raimondi, P., Jurgens, L. J., & Tinker, M. T. (2015). Evaluating potential conservation conflicts between two listed species: sea otters and black abalone. Ecology, 96(11), 3102–3108.

Raimondi, P., Wilson, C. M., Ambrose, R. E., Engle, J. M., & Minchinton, T. (2002). Continued declines of black abalone along the coast of California: are mass mortalities related to El Nino events? Marine Ecology Progress Series, 242, 143–152.

Richards, D. V., & Davis, G. E. (1993). Early warnings of modern population collapse in black abalone *Haliotis cracherodii*, Leach, 1814 at the California Channel Islands. Journal of Shellfish Research, 12, 189–194.

Robinson, J. A., Brown, C., Kim, B. Y., Lohmueller, K. E., & Wayne, R. K. (2018). Purging of strongly deleterious mutations explains long-term persistence and absence of inbreeding depression in island foxes. Current Biology: CB, 28(21), 3487–3494.e4.

Robinson, J. A., Kyriazis, C. C., Nigenda-Morales, S. F., Beichman, A. C., Rojas-Bracho, L., Robertson, K. M., Fontaine, M. C., Wayne, R. K., Lohmueller, K. E., Taylor, B. L., & Morin, P. A. (2022). The critically endangered vaquita is not doomed to extinction by inbreeding depression. Science, 376(6593), 635–639.

Rogers-Bennett, L. (2002). Estimating baseline abundances of abalone in California for restoration. CalCOFI Rep., 4.

Sanchez-Donoso, I., Ravagni, S., Rodríguez-Teijeiro, J. D., Christmas, M. J., Huang, Y., Maldonado-Linares, A., Puigcerver, M., Jiménez-Blasco, I., Andrade, P., Gonçalves, D., Friis, G., Roig, I., Webster, M. T., Leonard, J. A., & Vilà, C. (2022). Massive genome inversion drives coexistence of divergent morphs in common quails. Current Biology: CB, 32(2), 462–469.e6.

Santiago, E., Novo, I., Pardiñas, A. F., Saura, M., Wang, J., & Caballero, A. (2020). Recent demographic history inferred by high-resolution analysis of linkage disequilibrium. Molecular Biology and Evolution, 37(12), 3642–3653.

Sedlazeck, F. J., Rescheneder, P., Smolka, M., Fang, H., Nattestad, M., von Haeseler, A., & Schatz, M. C. (2018). Accurate detection of complex structural variations using single-molecule sequencing. Nature Methods, 15(6), 461–468.

Shaffer, H. B., Toffelmier, E., Corbett-Detig, R. B., Escalona, M., Erickson, B., Fiedler, P., Gold, M., Harrigan, R. J., Hodges, S., Luckau, T. K., Miller, C., Oliveira, D. R., Shaffer, K. E., Shapiro, B., Sork, V. L., & Wang, I. J. (2022). Landscape genomics to enable conservation actions: The California Conservation Genomics Project. The Journal of Heredity, 113(6), 577–588.

Sloan, N. A. (2003). Evidence of California-area abalone shell in Haida trade and culture. Can J Archaeol, 27(2), 273–286.

Subramanian, S. (2016). The effects of sample size on population genomic analyses--implications for the tests of neutrality. BMC Genomics, 17, 123.

Tajima, F. (1989). Statistical method for testing the neutral mutation hypothesis by DNA polymorphism. Genetics, 123(3), 585–595.

Teixeira, J. C., & Huber, C. D. (2021). The inflated significance of neutral genetic diversity in conservation genetics. Proceedings of the National Academy of Sciences of the United States of America, 118(10). 10.1073/pnas.2015096118

Terhorst, J., Kamm, J. A., & Song, Y. S. (2017). Robust and scalable inference of population history from hundreds of unphased whole genomes. Nature Genetics, 49(2), 303–309.

Tissot, B. N. (1988). Morphological variation along intertidal gradients in a population of black abalone *Haliotis cracherodii* Leach 1814. Journal of Experimental Marine Biology and Ecology, 117(1), 71–90.

Todesco, M., Bercovich, N., Kim, A., Imerovski, I., Owens, G. L., Dorado Ruiz, Ó., Holalu, S. V., Madilao, L. L., Jahani, M., Légaré, J.-S., Blackman, B. K., & Rieseberg, L. H. (2022). Genetic basis and dual adaptive role of floral pigmentation in sunflowers. eLife, 11. 10.7554/eLife.72072

VanBlaricom, G., Neuman, M., Butler, J. L., De Vogelaere, A., Gustafson, R. G., Mobley, C., Richards, D., Rumsey, S., & Taylor, B. L. (2009). Status review report for black abalone. National Marine Fisheries Service.

Van der Auwera, G. A., Carneiro, M. O., Hartl, C., Poplin, R., Del Angel, G., Levy-Moonshine, A., Jordan, T., Shakir, K., Roazen, D., Thibault, J., Banks, E., Garimella, K. V., Altshuler, D., Gabriel, S., & DePristo, M. A. (2013). From FastQ data to high confidence variant calls: the Genome Analysis Toolkit best practices pipeline. *Current Protocols in Bioinformatics / Editoral Board*, Andreas D. Baxevanis … [et Al.], 43(1110), 11.10.1–11.10.33.

Vileisis, A. (2020). Abalone: the remarkable history and uncertain future of California’s iconic shellfish. Oregon State University Press.

Villoutreix, R., Ayala, D., Joron, M., Gompert, Z., Feder, J. L., & Nosil, P. (2021). Inversion breakpoints and the evolution of supergenes. Molecular Ecology, 30(12), 2738–2755.

Wang, I. J., & Bradburd, G. S. (2014). Isolation by environment. Molecular Ecology, 23(23), 5649–5662.

Willi, Y., Kristensen, T. N., Sgrò, C. M., Weeks, A. R., Ørsted, M., & Hoffmann, A. A. (2022). Conservation genetics as a management tool: the five best-supported paradigms to assist the management of threatened species. Proceedings of the National Academy of Sciences of the United States of America, 119(1). 10.1073/pnas.2105076119

Zhang, C., Dong, S.-S., Xu, J.-Y., He, W.-M., & Yang, T.-L. (2019). PopLDdecay: a fast and effective tool for linkage disequilibrium decay analysis based on variant call format files. Bioinformatics, 35(10), 1786–1788.

